# Chromosome-level assemblies of *Amaranthus palmeri*, *Amaranthus retroflexus*, and *Amaranthus hybridus* allow for genomic comparisons and identification of a sex-determining region

**DOI:** 10.1101/2024.09.18.613719

**Authors:** Damilola A. Raiyemo, Jacob S. Montgomery, Luan Cutti, Fatemeh Abdollahi, Victor Llaca, Kevin Fengler, Alexander J. Lopez, Sarah Morran, Christopher A. Saski, David R. Nelson, Eric L. Patterson, Todd A. Gaines, Patrick J. Tranel

## Abstract

*Amaranthus palmeri* (Palmer amaranth), *A. retroflexus* (redroot pigweed), and *A. hybridus* (smooth pigweed) are troublesome weeds that are economically damaging to several cropping systems. Collectively referred to as ‘pigweeds’, these species are incredibly adaptive and have become successful competitors in diverse agricultural settings. Development of genomic resources for these species promises to facilitate the elucidation of the genetic basis of traits such as biotic and abiotic stress tolerance (e.g., herbicide resistance) and sex determination. Here, we sequenced and assembled chromosome-level genomes of these three pigweed weed species. By combining the haplotype-resolved assembly of *A. palmeri* with existing restriction site-associated DNA sequencing data, we identified a ∼2.84 Mb region on chromosome 3 of Hap1 that is male-specific and contains 37 genes. Transcriptomic analysis revealed two genes within the male-specific region, RESTORER OF FERTILITY 1 (*Rf1*) and TLC DOMAIN-CONTAINING PROTEIN (*TLC*), were upregulated in male individuals across the shoot apical meristem, the floral meristem, and mature flowers, indicating their potential involvement in sex determination in *A. palmeri*. In addition, we rigorously classified cytochrome P450 genes in all three pigweeds due to their involvement in non-target site herbicide resistance. Finally, we identified contiguous extrachromosomal circular DNA (eccDNA) in *A. palmeri,* a critical component of glyphosate resistance in this species. The findings of this study advance our understanding of sex determination in *A. palmeri* and provide genomic resources for elucidating the genetic basis and evolutionary origins of adaptive traits within the genus.

## Introduction

*Amaranthus palmeri* S. Watson (Palmer amaranth), *A. retroflexus* L. (redroot pigweed), and *A. hybridus* L. (smooth pigweed) are troublesome summer annual weeds of several cropping systems (Sellers et al. 2003). While *A. retroflexus* and *A. hybridus* are monoecious and primarily self-pollinated, *A. palmeri* is a dioecious (i.e., with separate male and female plants), obligate outcrossing species that has now spread beyond its native origins in the American Southwest to many parts of the USA and the world (Steckel 2007; Trucco et al. 2007). *A. palmeri* has now been reported in 45 countries, and climate models indicate that it will expand to most temperate and subtropical areas of the USA, Africa, Australia, South America, Eastern Asia, and Europe by 2050 (Kistner and Hatfield 2018; Roberts and Florentine 2022). Several studies have documented significant yield losses that could occur from interference of weedy amaranths (collectively referred to as ‘pigweeds’) with diverse crops such as corn, soybean, or cotton (Bensch et al. 2003; Hager et al. 2002; Morgan et al. 2001; Massinga et al. 2001; Manalil et al. 2017).

The notoriety of the pigweeds as damaging to agricultural crops has spurred the exploration of several options for their control (e.g., cover crops, herbicide rotations, mechanical seed destruction, flood irrigation management, tillage, and grazing). A meiotic gene drive of a sex-determining genetic factor has been proposed as a radically new method for suppression of dioecious weed species such as *A. palmeri* (Barrett et al. 2019; Legros et al. 2021; Rode et al. 2019; Neve 2018). To utilize such a weed control strategy, however, requires understanding the mechanism of sex determination in such a species. Previous studies on sex determination in amaranths proposed that males are the heterogametic sex with an XY system, which was subsequently confirmed using restriction-site associated DNA (RAD) sequencing (Murray 1940; Grant 1959; Montgomery et al. 2019; Neves et al. 2020). A ∼1.3 Mb male-specific Y region (MSY) that contained 121 gene models was identified in a draft genome assembly of a male *A. palmeri* plant (Montgomery et al. 2020, 2021). The draft assembly was also combined with short reads of *A. watsonii* to detect the male-specificity of a copy of a gene encoding a pentatricopeptide repeat-containing protein (PPR) within the MSY region on scaffold 20 (Raiyemo et al. 2023). Similarly, transcriptomic analyses between several tissue types of male and female individuals revealed the upregulation of *PPR247* in males, a gene within the MSY region (Bobadilla et al. 2023). The male-specific expression of this gene across three tissue types led Bobadilla et al. (2023) to hypothesize a single-gene model of sex determination for *A. palmeri*. Despite the advances above, the genomic landscape of sex chromosomes and how they evolved for the dioecious *Amaranthus* species are still poorly understood. For example, inversions, centromeric location of sex-determining regions or hemizygosity have been reported as possible routes of recombination suppression that lead to the evolution of sex chromosomes in some dioecious species (Carey et al. 2022). In fact, we recently reported the identification of a sex-determining region in the closely related *Amaranthus tuberculatus*, near a chromosomal fusion and large inversion event (Raiyemo et al., 2024).

Evolution and spread of herbicide resistance in pigweeds is now a major concern for crop production. Non-target-site herbicide resistance (NTSR), specifically metabolism-based resistance, is particularly alarming due to its genetic complexity and the ability of one resistance allele to give cross resistance to diverse herbicides (Yu and Powles 2014; Rigon et al. 2020). Cytochrome P450 (P450) genes are involved in enhanced herbicide metabolism, and are known to mediate the detoxification of herbicides by adding a functional group to the herbicide molecule via oxidation, reduction or hydrolysis, thus rendering the herbicide less effective (Gaines et al. 2020; Dimaano and Iwakami 2021). In fact, P450s have been implicated in metabolism-based herbicide resistance of some *A. tuberculatus* and *A. palmeri* biotypes (Oliveira et al. 2018; Varanasi et al. 2018; De Figueiredo et al. 2022; Rigon et al. 2023). Understanding the structure-activity relationship between CYP enzymes and the herbicides they metabolize via biochemical modeling could help predict herbicide classes that could be degraded even before they are synthesized (Gaines et al. 2020; Bobadilla and Tranel 2024). Development of genome assemblies and careful annotation of these P450 genes for weed species of interest provides a library of herbicide metabolism genes for systematic approaches to such investigations.

Furthermore, the identification of gene duplication of *5-enolpyruvylshikimate 3-phosphate synthase* (*EPSPS*) on extrachromosomal circular DNA (eccDNA) conferring resistance to the herbicide glyphosate in *A. palmeri* was a surprising and landmark discovery for weed genomics (Gaines et al. 2010; Koo et al. 2018; Molin et al. 2020b). Since this discovery, eccDNAs have been found in many other species, and variation of eccDNAs within and among species has become an open area of research for *A. palmeri* and other species (Camposano et al. 2022; Fu et al. 2023). It has been suggested that the eccDNAs that contain *EPSPS* in *A. palmeri* are diverse with multiple evolutionary events (Camposano et al. 2022), but the source of eccDNAs are not well understood. A better understanding of the evolutionary origins of eccDNAs and how they amplify genes will shed light on this surprising mechanism of gene duplication and determine the necessity of resistance containment protocols.

Towards furthering our understanding of the biology of weed species including pigweeds, the International Weed Genomics Consortium (IWGC) was initiated to provide genomic resources for the science community (Montgomery et al. 2024). As part of this effort, we produced chromosome-level genome assemblies of *A. palmeri* (male; haplotype-resolved), *A. retroflexus*, and *A. hybridus* individuals utilizing PacBio long-read sequencing, Hi-C scaffolding, and Bionano optical mapping. Utilizing these genome assemblies, we identified a contiguous male-specific region and candidate genes for sex determination in *A. palmeri*, characterized the P450 gene family in the three pigweeds, and identified the conservation of eccDNA sequence across diverse accessions of *A. palmeri*. The genome assemblies reported here along with existing *Amaranthus* genomes promises to improve weed management through development of new weed control technologies as well as crop improvement through identification of the genetic basis of stress tolerance in these prolific species.

## Results

### Genome assembly, annotation and repeat analyses

Although the assemblies contained a number of unresolved gaps corresponding to AT-rich regions, analyses of *A. palmeri* haplome 1 (Hap1), *A. palmeri* haplome 2 (Hap2), *A. retroflexus* and *A. hybridus* genomes using BUSCO and LTR assembly index (LAI) revealed highly complete assemblies for all species (Table 1). The total number of predicted protein-coding genes for the assemblies ranged from 22,771 genes for *A. hybridus* to 27,377 genes for *A. retroflexus* (Table 1; Supplemental Table S1 and S2). Analysis of repetitive elements revealed over half of each genome assembly was made up of transposable elements with LTR/*Copia* elements being the most abundant (∼10% of each genome) (Supplemental Table S3). Analysis of centromeric repeats indicates chromosomes have varying types of centromeres. For example, chromosomes 1 and 5 appear telomeric while chromosomes 2 and 3 appear metacentric or submetacentric for *A. palmeri* (Supplemental Table S4). A search of simple telomeric repeat against the assemblies revealed telomeric repeat sequences at ∼75% of chromosome ends for *A. retroflexus* and both haplotypes of *A. palmeri*, but just 25% of chromosome ends for *A. hybridus* (Supplemental Table S5). Localization of genomic features depicted in Figure 1 indicates that regions abundant in gene content are poor in LTR retrotransposons (LTR-RT), and vice-versa.

**Table 1.** Comparison of assembly statistics between the two haplomes of *A. palmeri*, *A. retroflexus*, and *A. hybridus* genome assemblies.

| Genome characteristics | *A. palmeri* |  | *A. retroflexus* | *A. hybridus* |
| --- | --- | --- | --- | --- |
|  | Haplome 1 | Haplome 2 |  |  |
| Assembly size (Mbp) | 384.1 | 364.5 | 446.79 | 437.99 |
| Scaffold N50 (Mbp) | 23.59 | 22.01 | 24.31 | 27.31 |
| Scaffold L50 | 8 | 8 | 8 | 7 |
| GC content (%) | 33.4 | 33.4 | 33.0 | 33.0 |
| Complete BUSCO (%) | 97.7 | 97.3 | 97.3 | 97.3 |
| Size of Ns (Mbp) | 19.14 | 7.02 | 21.64 | 40.79 |
| LTR assembly index (LAI) | 18.25 | 21.85 | 10.78 | 15.29 |
| Protein-coding genes | 24,873 | 24,791 | 27,377 | 22,771 |
| Mean gene length (bp) | 4,637 | 4,633 | 4,623 | 4,874 |
| Mean CDS length (bp) | 1,178 | 1,174 | 1,092 | 1,184 |
| Mean exon length (bp) | 273 | 272 | 298 | 270 |
| Mean exon per gene | 5.5 | 5.5 | 5.2 | 5.5 |
| Number of tRNA | 1,480 | 1,508 | 1,499 | 1,535 |
| Total annotated genes | 26,353 | 26,299 | 28,876 | 24,306 |

**Figure 1.**
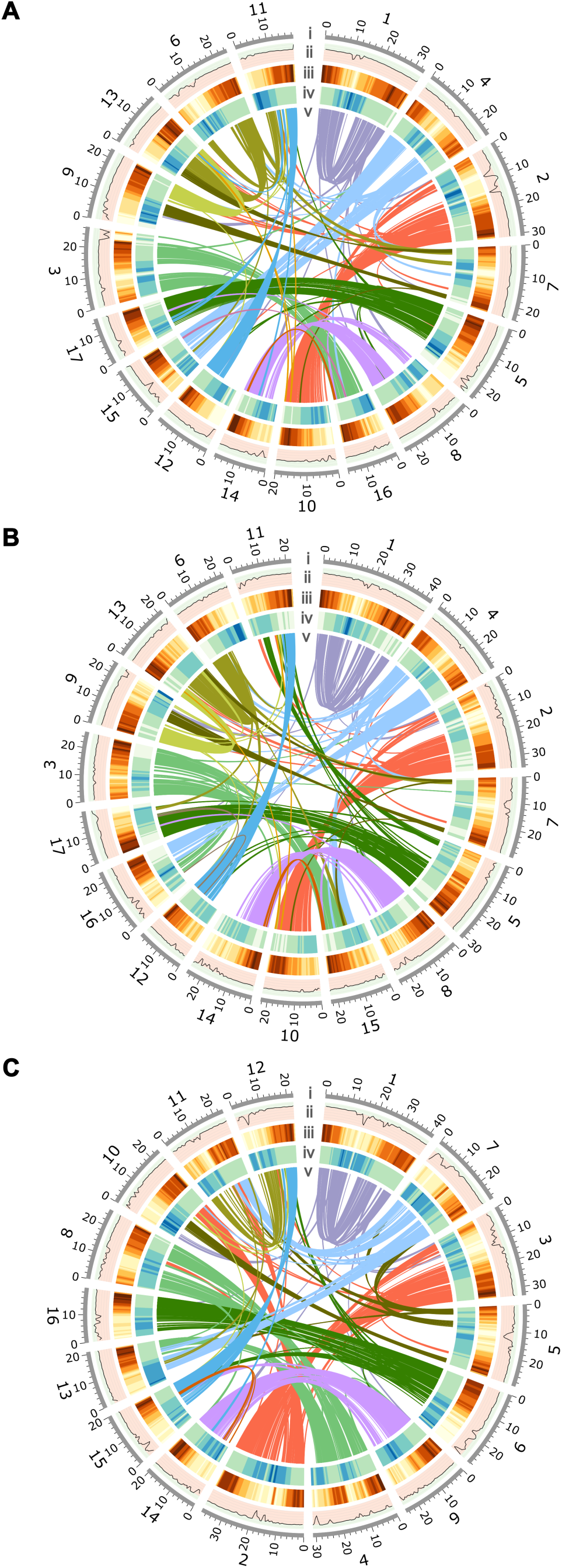
Genomic features. (*A*) *A. palmeri* Hap1. (*B*) *A. retroflexus*. (*C*) *A. hybridus* assemblies. Circos plot depicts i) chromosome number and length (Mb), ii) GC content along the chromosomes, with peaks in light green area representing GC content greater than the median and peaks in light red area representing GC content less than the median (median GC contents: ∼ 0.32), iii) gene density across the chromosomes, with brown representing gene-rich regions and yellow representing gene-poor regions, iv) Long terminal repeats (LTR) density along chromosomes, with blue representing LTR-rich regions and green representing LTR-poor regions, v) inner ribbons represent duplicated genes. Window size of 1 Mb and step size of 500 Kbp for ii-iv.

### Interspecific comparisons of genome structure

Analysis of structural rearrangements among the three *Amaranthus* genome assemblies from this study, *A. tuberculatus*, and three monoecious amaranths indicated a highly conserved gene order, except for a few chromosomes (Fig. 2). For instance, chromosome 4 in *A. hypochondriacus*, *A. hybridus*, and *A. cruentus* appears to have been derived from the fusion of two ancestral chromosomes (Fig. 2). It is worth noting that some genes on chromosome 1 of the three genome assemblies appear to have paralogs (i.e., duplications) on chromosome 1 (Fig. 1). This pattern of gene duplication on chromosome 1 was earlier observed in *A. hypochondriacus* (Lightfoot et al. 2017) and *A. cruentus* (Ma et al. 2021), and thus seems to be conserved across species in the subgenera *Acnida* and *Amaranthus*. Chromosome 1 (or chromosome 2 in *A. tuberculatus*) in species within the two subgenera, however, appears to have originated from the fusion of parts of chromosomes 1 and 2 in *A. tricolor*, which belongs to the subgenus *Albersia* (Fig. 2).

**Figure 2.**
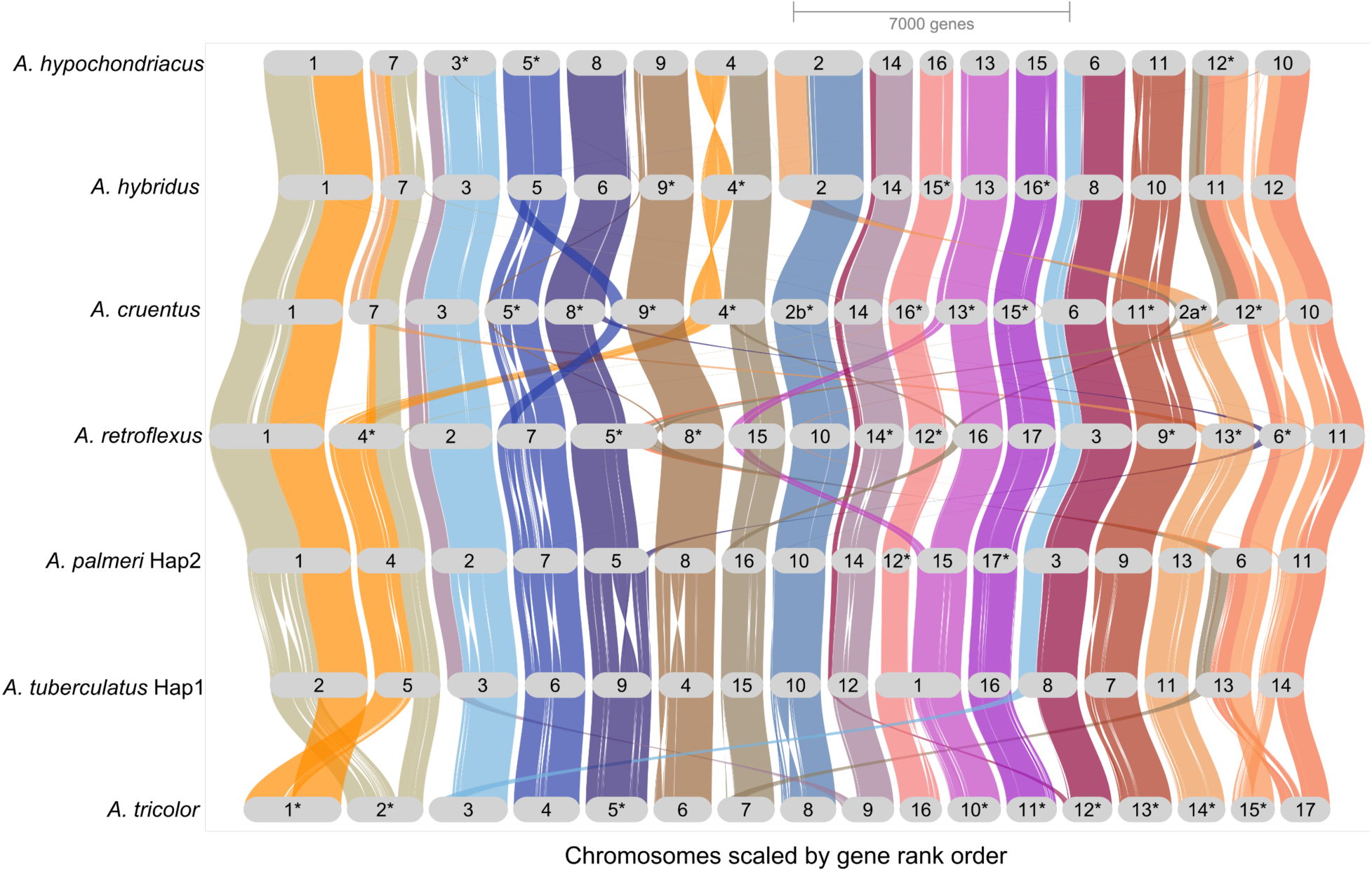
Synteny plot between the haplotype assemblies of four previously reported chromosome-level *Amaranthus* spp. assemblies and three newly assembled ones (*A. palmeri*, *A. retroflexus*, and *A. hybridus*). Asterisks (*) indicate chromosomes that were manually inverted to keep the gene order consistent with *A. tuberculatus*. Species are ordered to reflect phylogenetic relationships from STAG in OrthoFinder, which also corresponds to the species tree in Wang et al. (2023).

### Identification of the sex-determining region (SDR) in *A. palmeri*

Individuals with more than 60% missing data were removed from a previously produced RAD-seq dataset, which retained 52 female samples and 53 male samples from across six geographically diverse accessions. The filtering improved the quality of the data (i.e., 24.57% missing data across the 105 individuals). The individuals were then mapped to Hap1 of the genome assembly, and 234,066 variants retained across the individuals were used for genome-wide association (GWA) analysis. The analysis revealed four significant SNPs above the Bonferroni threshold that are associated with sex; two on chromosome 3 (24,609,596 bp, *P* = 1.7481e-20; 24,595,358 bp, *P* = 6.8453e-17), and one each on chromosome 4 (13,423,105 bp, *P* = 1.0254e-08), and chromosome 16 (7,114,501 bp, *P* = 1.0704e-17) (Fig. 3A). All significant SNPs were within intergenic regions, and there was no evidence of systematic bias in the GWA analysis based on the QQ plot (Fig. 3B). Further analysis of genetic differentiation between females and males using fixation index (*F_ST_*) also points to the end of chromosome 3 as being highly differentiated between female and male individuals of *A. palmeri* (Figs. 3C, 3D). Given the location of the sex-determining region, we therefore designate Hap1 as the assembly containing the Y chromosome while Hap2 contains the X chromosome.

Analysis of synteny between the haplotype assemblies indicated a mostly one-to-one relationship in gene content between the haplotypes (Supplemental Figure S1A and S1B). Identification of syntenic orthologs revealed 21,421 gene pairs are reciprocal best hit matches between the two haplotypes. A region on chromosome 3 between 22,994,134 – 25,836,006 (∼ 2.84 Mb) is hemizygous and present near the end of Hap1 but absent from Hap2. In the absence of this ∼2.84 Mb region, Hap1 and Hap2 were highly syntenic (Figure 3E). Further analysis revealed 1,857,183 bp (65.35%) of this hemizygous region of Hap1 is made up of Ns, suggesting large stretches of simple repeats that were not assembled. While the sequence is not represented, the length of gap is known because BioNano maps estimate the physical distance between markers at well-assembled loci. There are 37 genes within this hemizygous region out of 1,620 protein-coding genes on chromosome 3 of Hap1 (Supplemental Table S6). Analysis of structural rearrangements indicated inversions were also present on chromosome 3. They were, however, less than 260 Kbp; shorter than other inversions on chromosomes 4, 7, 14, or 17 that were greater than 1 Mb (Supplemental Table S7).

Query of a primer set (PAMS-940), used in Montgomery et al. (2019) as a male-specific marker, against both haplotype assemblies of *A. palmeri* revealed a perfect match to chromosome 3 of Hap1 with a predicted amplicon size of 51 bp (24,464,780 – 24,464,831 bp) (Fig. 3E). This size is consistent with the amplified product size reported in Montgomery et al. (2019). The forward and reverse primer sequence for JM940 from Montgomery et al. (2021) also matched to chromosome 3 (24,755,051 - 24,755,052), but two single nucleotide polymorphisms exist between the forward primer and the genomic sequence. Sequence alignment of the previous draft *A. palmeri* male assembly to both haplotype assemblies identified large structural differences between assemblies (Supplemental Figs. S2A and S2B). This variation could be due to real differences in genome structure, but also could be due to errors in assembly or scaffolding. Scaffolds 5 (previously proposed as containing the pseudoautosomal region) and 20 (containing a previously proposed male-specific region) both aligned to chromosome 3 in the two haplotype assemblies. As expected, the region identified as male-specific on scaffold 20 only aligned to the hemizygous region on chromosome 3 of Hap1 (Supplemental Figs. S2A–S2F). Reciprocal best hit searches between genes on scaffold 20 of the draft assembly and those in the haplotype assemblies revealed 41 and 37 genes had matches on chromosome 3 of Hap1 and Hap2 assemblies, respectively (Supplemental Table S8). However, only four genes (coding for three unknown proteins and HEADING DATE 3A) that matched between the draft assembly and Hap1 were within the male-specific region in Hap1 (Supplemental Table S8). Taken together, we consider chromosome 3 as the likely sex chromosome candidate in our assembly and the ∼2.84 Mb hemizygous region as the likely sex-determining region.

**Figure 3.**
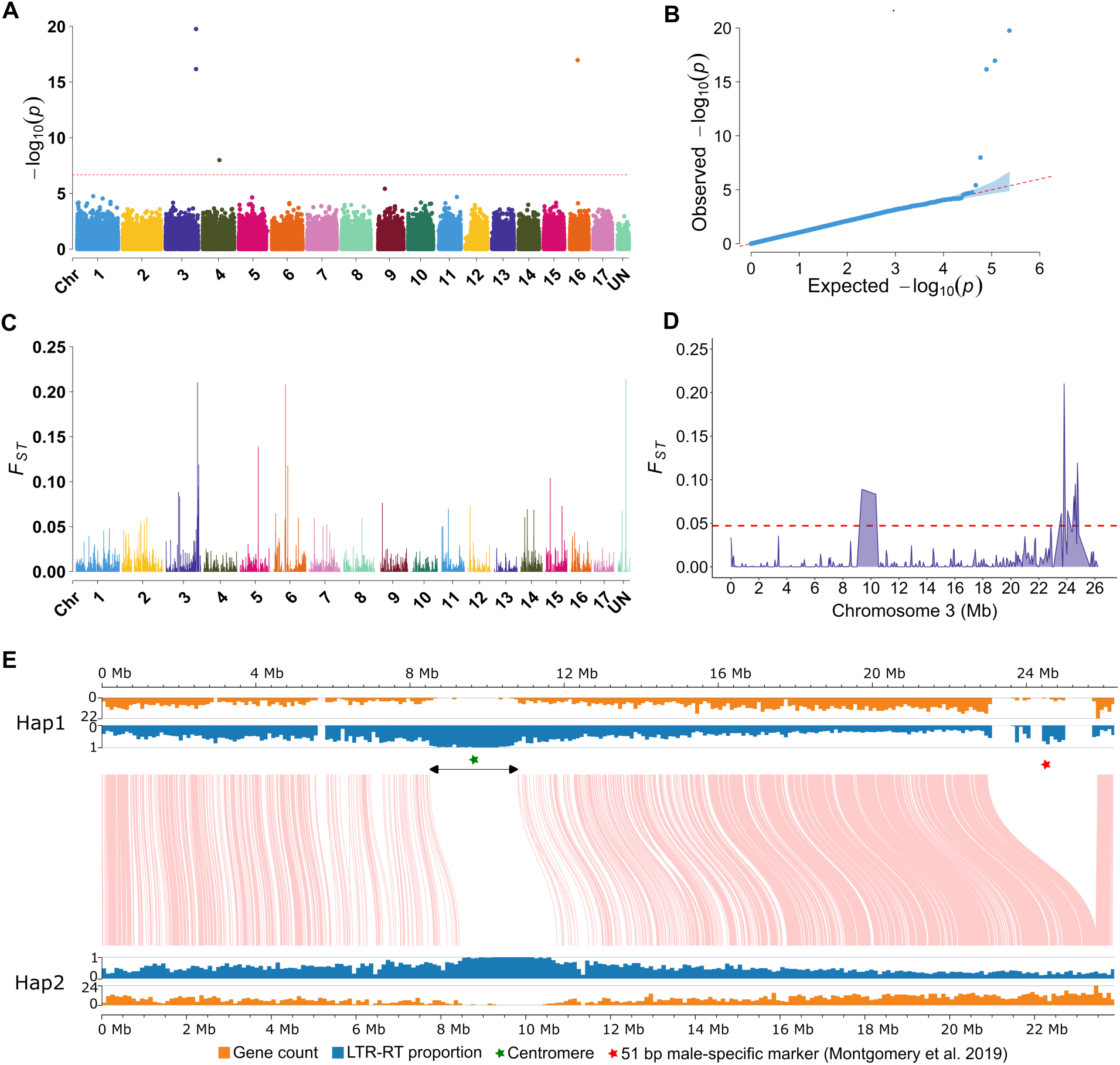
Identification of *A. palmeri* sex-determining region on chromosome 3. (*A*) Manhattan plot of GWA analysis using RAD-seq data from 52 females and 53 males. The dashed red line indicates Bonferroni threshold of –log_10_(*P*) = 6.6704. (*B*) Quantile-quantile (QQ) plot of the GWA analysis. (*C*) Fixation index (*F_ST_)* between females and males across all 17 chromosomes (window size 100 Kbp; step size 50 Kbp). (*D*) *F_ST_* between females and males for chromosome 3 with dashed red line representing the top 1% threshold at 0.0472 (window size 100 Kbp; step size 50 Kbp). (*E*) Synteny plot showing collinear regions on chromosome 3. The green asterisk represents the centromere while the red asterisk represents a previously reported 51 bp male-specific marker (Montgomery et al. 2019) that matched to a region on Hap1. Gene count and LTR-RT proportion were calculated based on 100 Kbp non-overlapping windows.

### Transcriptome profiling, gene ontology term enrichment, and coverage analyses

A previous mRNA dataset reported in Bobadilla et al. (2023) was mapped to the Hap1 assembly of *A. palmeri*. Uniquely mapped reads ranged from 76.40 – 91.61% across male and female samples for shoot apical meristem, floral meristem, and mature flower tissues. Out of the 24,873 annotated Hap1 protein-coding genes, 20,046 genes were retained for differential expression (DE) analysis after filtering and trimmed mean of M values (TMM) normalization (Supplemental Figs. S3A and S3B). Eight genes were differentially expressed between males and females in the shoot apical meristem, while 29 genes were differentially expressed for floral meristem, and 2,595 genes for mature flower (Supplemental Figs. S3A and S3B). Only five genes (encoding a serine/threonine-protein kinase EDR1-like protein, two Rf1 proteins, an unknown protein, and a TLC domain-containing protein) were differentially expressed between male and female individuals across the three tissue types, and all mapped to chromosome 3 (Supplemental Tables S9 – S11). A search of the unknown protein to NCBI nonredundant protein database using BLASTP revealed 71% homology to a wall-associated receptor kinase-like protein in several species. Two of these five genes, encoding either an Rf1 protein (a pentatricopeptide repeat-containing restorer of male fertility gene) (AmaPaChr03Ag063970) or a TLC domain-containing protein (AmaPaChr03Ag063980), are within the MSY region on chromosome 3 (Supplemental Tables S9 – S11). Gene ontology (GO) term enrichment analysis, with mature flower DEGs selected based on a false discovery rate threshold of p < 0.05 and Log_2_FoldChange > 1.2, revealed the list of DEGs was predictably enriched for genes involved in pollen and anther development (Supplemental Fig. S3C, Supplemental Table S12 – S15).

To determine the conservation of these two genes with male-specific expression in our predicted SDR, we mapped genomic sequence data of *Amaranthus watsonii*, a species closely related to *A. palmeri* (Raiyemo et al. 2023; Raiyemo and Tranel 2023), to the Hap1 genome assembly of *A. palmeri*. Coverage analysis revealed that while the *TLC domain-containing* gene that was upregulated in males and within the MSY region has male-biased coverage in *A. watsonii* (Fig. 4A), the *Rf1* gene that was also upregulated in males and within the MSY region appears to be male-specific in *A. watsonii* (Fig. 4B). After filtering to retain only uniquely mapped reads, the *TLC domain-containing* gene no longer had any reads aligning from male or female *A. watsonii* plants while the *Rf1* gene still had male-specific coverage for the longest exon (Supplemental Fig. S4). Similarly, a gene encoding a different pentatricopeptide repeat-containing protein (AmaPaChr03gA063950), although not differentially expressed between males and females, was male-specific in *A. watsonii* even after filtering for only uniquely mapped reads (Supplemental Fig. S5). Other genes of unknown function within the MSY of *A. palmeri* also showed male-enriched coverages in *A. watsonii* (Supplemental Figs. S5, S6). Given their location in the SDR region, male-specific expression, and conservation across species, we hypothesize that *Rf1* and *TLC* are necessary for maleness and/or suppression of femaleness in *A. palmeri* and *A. watsonii*.

**Figure 4.**
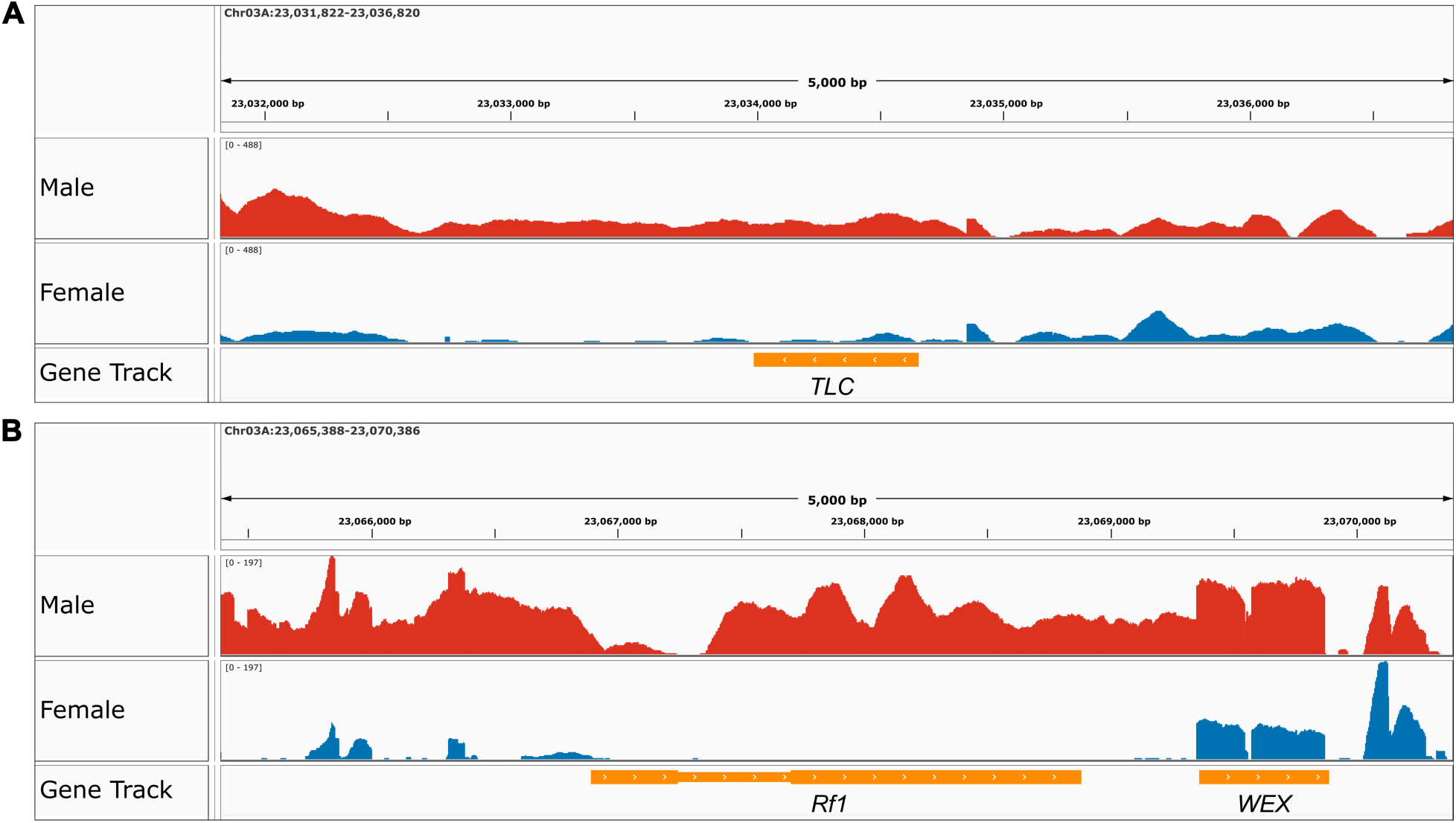
Male-specific gene coverage analysis. Reads of male and female *A. watsonii* mapped to *A. palmeri* Hap1 assembly and visualized in IGV. (*A*) TLC domain-containing protein (AmaPaChr03Ag063980). (*B*) Rf1 protein (AmaPaChr03Ag063970).

### Identification and comparative analyses of *Amaranthus* cytochrome P450 gene superfamily

A total of 184, 214, and 228 cytochrome P450 (CYP) gene models were identified in *A. palmeri, A. retroflexus* and *A. hybridus*, respectively and named according to the Standardized Cytochrome Nomenclature Committee (http://drnelson.uthsc.edu/CytochromeP450.html) (Supplemental Tables S16–S18). Among these gene models, 130 from *A. palmeri*, 163 from *A. retroflexus*, and 152 from *A. hybridus* were determined to be full-length (ranging from 350 to 700 amino acids) while the rest (less than 350 amino acids) were designated as fragments and discarded from the analysis (Supplemental Tables S16). The full-length protein sequences for all species are provided in Supplemental Table S19.

A neighbor-joining (NJ) tree was constructed from protein sequence to investigate the evolutionary relationship among the full-length CYP genes (Supplemental Figs. 7A, 7B). Genes were thus categorized into two groups: A-type, with a single clan (71), and non-A-type with multiple clans including 710, 85, 711, 86, 97, 72, 727, and 74. Some clans (74, 97, 710, 711, 727) contained only one gene family each. In contrast, other clans, namely 71, 72, 85, and 86, included multiple families (Fig. 7A, 7B). In total, 50% of CYP genes in *A. palmeri* (65/130), 45.3% in *A. retroflexus* (74/163), and 44.5% in *A. hybridus* (81/182) belonged to clan 71, which emerged as the largest clan and includes 17 families (CYP71, CYP81-82, CYP92-93, CYP75-79, CYP84, CYP701, CYP703, CYP706, CYP89, CYP98, and CYP736). Here, clans are higher-level groupings of P450 genes that contain several families, whereas a CYP family is a more specific classification within a clan that is based on higher sequence identity (at least 40%) and closer evolutionary relationships.

Collinearity analysis was also conducted to explore the evolutionary relationships among the CYP gene family of the three species. We found a total of 106 collinear gene pairs between *A. palmeri*, and *A. hybridus* (82% of *A. palmeri* CYPs), as well as 95 collinear gene pairs between *A. palmeri* and *A.retrofloxus* (73% of *A. palmeri* CYPs) (Fig. 7C). These results highlight the significant evolutionary conservation present within the CYP genes among *A. palmeri, A. retroflexus* and *A. hybridus*.

### Origin of extrachromosomal circular DNA (eccDNA) and gene amplification in *A. palmeri*

To understand the genomic origins of eccDNA containing the gene *EPSPS* and conferring glyphosate resistance, the complete assembly of a 399 Kbp eccDNA molecule reported in Molin et al. (2020) was aligned to the reference genome (Hap1) of *A. palmeri*, which was obtained from a glyphosate-susceptible individual. Relatively stringent filtering (map quality = 60 and alignment length >1000 bp) revealed few syntenic regions between the canonical *EPSPS*-containing eccDNA molecule and the genome (Supplemental Figure S8A). The longest stretch of sequence similarity was the region containing *EPSPS* (∼10 Kbp). However, relaxing the filtration of alignments (95% similarity, minimum alignment length>30 bp) resulted in many matches across the genome (Supplemental Figure S8B).

To understand the evolutionary history of *EPSPS* amplified via eccDNAs, geographically diverse glyphosate-susceptible and glyphosate-resistant samples of *A. palmeri* were sequenced and their genomes assembled. When the assemblies of the glyphosate-sensitive individuals were aligned to the canonical eccDNA molecule, the assemblies had few, short alignments compared to glyphosate-resistant individuals (Supplemental Tables S20-27). This suggests the glyphosate-sensitive samples do not have an “empty” version of the eccDNA molecule that is simply missing *EPSPS*. In contrast, glyphosate-resistant samples had contigs that aligned across the entirety of the canonical eccDNA molecule, in line with findings from Molin et al. (2020a).

After collapsing redundant contigs, full *EPSPS* gene models were only found on one contig for each glyphosate-susceptible sample. Assemblies of glyphosate-resistant samples retained between 4 and 13 contigs containing *EPSPS* gene models. Alignment of these *EPSPS*-containing contigs to the canonical *EPSPS*-containing eccDNA sequence showed complete synteny between the contigs and the eccDNA sequence (Fig. 5A). While most contigs were only fragments of the canonical eccDNA, a 395 Kbp circular contig containing *EPSPS* was recovered from the assembly of a glyphosate-resistant plant from Georgia. Though its length is similar to the eccDNA molecule reported by Molin et al. (2020), there are several large insertions (up to 3 Kbp) and deletions (up to 8 Kbp) in the *de novo* assembled version and very few single nucleotide polymorphisms. These insertions and deletions within the eccDNA sequence are mainly found in regions of simple repeats and could be due to sequencing artifacts. However, the insertions and deletions are conserved across the four diverse glyphosate-resistant samples, suggesting that they are likely not sequencing artifacts, but widespread among resistant samples. Genomic versions of *EPSPS* were found in all samples on contigs that did not have synteny with the canonical eccDNA molecule, but these were determined to be assemblies of the genomic locus containing *EPSPS* on chromosome 7.

A phylogeny of *EPSPS* gene sequences from all sequenced individuals and several outgroup species clustered all *A. palmeri* sequences together, indicating *EPSPS* from an *A. palmeri* plant was inserted into the eccDNA molecule (Fig. 5B). Koo et al. (2023) showed that interspecific hybridization can transfer this eccDNA to sexually compatible species of *Amaranthus*, but this result confirms that it originally evolved in *A. palmeri*. Further, Figure 5B shows that all eccDNA versions of *EPSPS* group very tightly together, suggesting insertion of *EPSPS* into the eccDNA occurred only once, evolutionarily recently, followed by rapid geographic spreading through gene flow.

**Figure 5.**
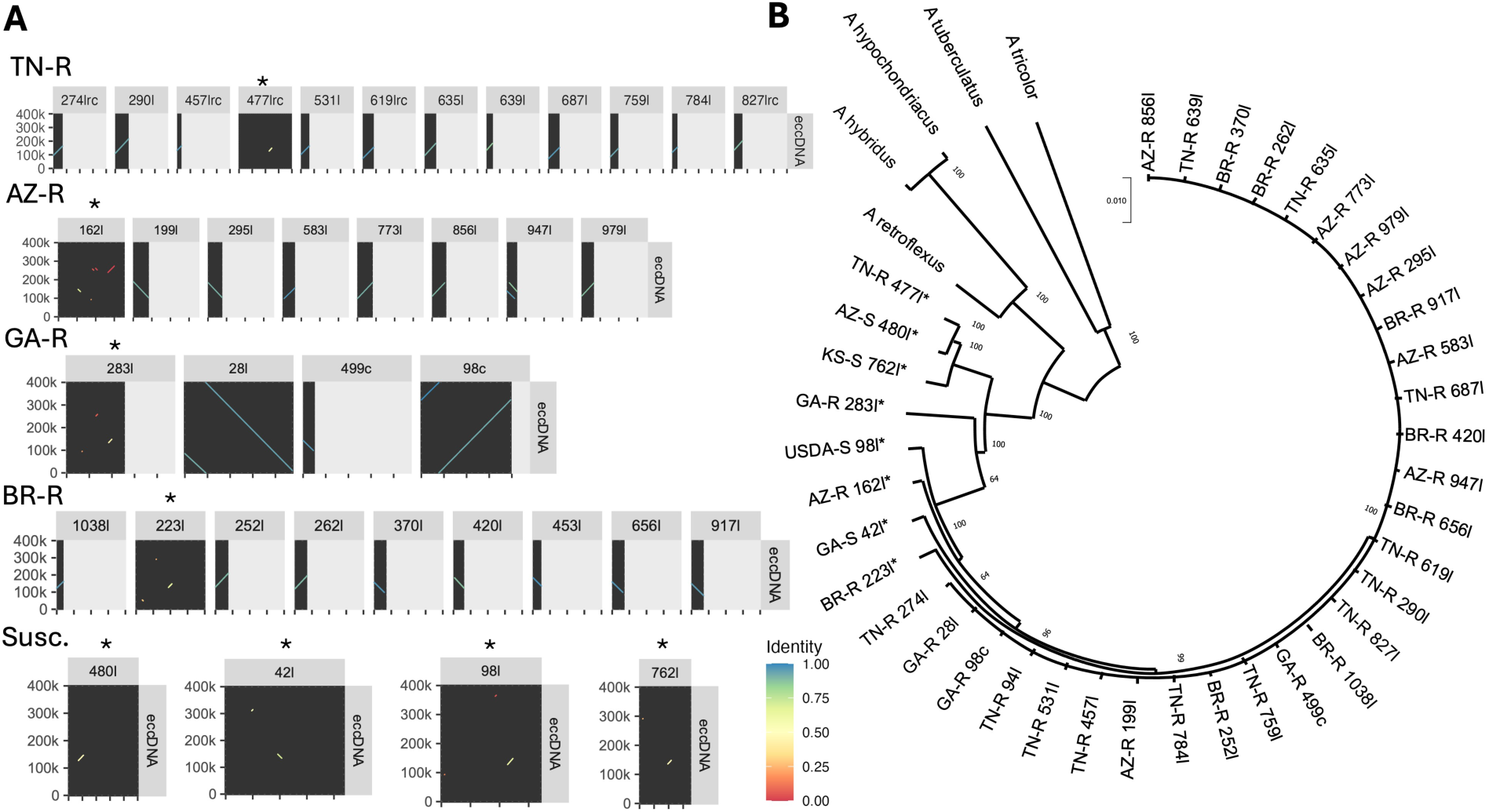
Comparison of extra-chromosomal circular DNA (eccDNA) containing *EPSPS* gene models from *A. palmeri*. (*A*) Dotplots showing synteny between the canonical eccDNA molecule reported by Molin et al. (2020) and assembled contigs containing *EPSPS* gene models (located near 140Kbp on the eccDNA) from diverse *A. palmeri* accessions. Each row contains contigs from a different individual, except the last row contains the contigs from all four glyphosate-susceptible (Susc.) samples. (*B*) Maximum likelihood phylogenetic tree estimating evolution of *EPSPS* genes within contigs of *de novo* assemblies of each resequenced *A. palmeri* accession. *EPSPS* sequences from other *Amaranthus* species were used as out groups. Values at each branch indicate the number of bootstrap iterations that generated that branch (n=100). Names of each branch indicate species or *A. palmeri* accession, contig number, and whether that contig was circularized or linear (l=linear, c=circular, rc=reverse complement). * indicates contigs that are part of the nuclear genome. Arizona-resistant (AZ-R), Arizona-susceptible (AZ-S), Brazil-resistant (BR-R), Georgia-resistant (GA-R), Georgia-susceptible (GA-S), USDA-susceptible (USDA-S), Tennessee-resistant (TN-R), Kansas-susceptible (KS-S).

## Discussion

We present the first haplotype-resolved chromosome-level assemblies of *A. palmeri*, *A. retroflexus*, and *A. hybridus*; three species that have become troublesome weeds in numerous agricultural systems around the world. Genome assembly identified chromosomal rearrangements that led to diversity in chromosome numbers across the genus. Our analysis identified an approximately 2.84 Mb region near the end of chromosome 3 on *A. palmeri* Hap1 that is male-specific, which is longer than the 1.3 Mb region that was previously identified as male-specific for the species (Montgomery et al. 2021). The difference in sizes is likely due to the inclusion of optical mapping in this study, which allowed us to represent unassembled regions as stretches of N’s. With 65.35% of the male-specific or Hap1-linked region made up of N-gaps, it is possible that our assembly is still missing male-specific sequences within the region. Considering that some of the gaps are flanked by AT repeats or other combinations of simple sequence repeats, it is possible the sequences within the gaps are difficult-to-assemble repeats that pose assembly challenges. Nevertheless, we identified key genes (encoding Rf1 and TLC domain-containing protein) that are likely involved in sex determination. While the function(s) of TLC domain-containing proteins are unclear in plants, Rf1 proteins are well-documented as restorers of fertility in some crops including sorghum and wheat (Klein et al. 2005; Melonek et al. 2021). More so, over half of cloned restorer (*Rf*) genes encode pentatricopeptide repeat-containing (PPR) proteins with roles in restoring normal pollen production to plants carrying a cytoplasmic male sterility locus (Gaborieau et al. 2016; Uyttewaal et al. 2008; Chen and Liu 2014).

Bobadilla et al. (2023) reported three genes, *PPR247*, *ACD6*, and *WEX*, as likely candidates involved in sex determination in *A. palmeri*. They proposed that the presence of *PPR247* within the MSY region of *A. palmeri* results in the post-transcriptional silencing of *ACD6* and *WEX*, thus enabling the formation or development of male reproductive organs. In our study, we identified an *Rf1* gene that shares homology with their identified *PPR* gene, but phylogenetic clustering places these genes on different branches (Supplemental Fig. S9). The other genes proposed by Bobadilla et al. (2023) fall outside of the male-specific region and are therefore not likely to be male determinants, although they could play a role in sex expression.

Recently, Wu et al. (2023) showed that while staminate flowers in *A. palmeri* initiate both carpel and stamen primordia at an early stage, the carpel remains undeveloped, and individuals become functionally male, whereas pistillate flowers initiate and develop only carpel primordium. Given that both androecium and gynoecium are initiated in staminate flowers, but only functional androecium develops, it is plausible that the expression of a dominant activator of maleness (*M*) at stage 4/5 of floral organogenesis (Wu et al. 2023) could result in the initiation and establishment of stamen primordia, whereas the expression of a dominant suppressor of female organs (*supF*) could result in the arrest of the initial gynoecium primordia at stage 7. It is therefore possible that the *Rf1* gene within the MSY region of *A. palmeri* is the male-promoting factor, and its presence and expression results in the formation of male reproductive organs. However, whether the *Rf1* gene also acts as the suppressor of female function is not clear. Knocking out the *Rf1* gene should result in male-to-female conversion if it acts as both the male activator and female suppressor but should cause a male-to-sterile conversion if a separate gene acts as the suppressor of female function.

The similarity between chromosome 3 of Hap1 and Hap2 assemblies besides the MSY region in Hap1 suggests that hemizygosity (i.e., presence of the MSY region near the end of chromosome 3 in Hap1 but absence in Hap2) is responsible for the recombination suppression between the two haplotypes. In contrast, large inversions within the sex-determining region on chromosome 1 of *A. tuberculatus* were identified and might mediate recombination suppression in that species (Raiyemo et al. 2024). Similar to our findings for *A. palmeri*, reduced recombination due to hemizygosity has been reported in garden asparagus (Harkess et al. 2020). The lack of one-to-one synteny between genes in the sex-determining regions of *A. palmeri* and *A. tuberculatus* further supports our previous conclusions on independent evolution of sex-determining regions in both species (Figure 2). However, the similarity between the gene ontology terms enriched for floral development in both species suggests that the downstream genes or pathway recruited in sex expression are similar. For instance, the most significantly overrepresented terms for male versus female differentially expressed genes in both species were pollen tube growth and regulation of pollen tube growth.

Beyond sex determination, comparative analyses also provided insights into cytochrome P450s and integration of *EPSPS* into eccDNA, within *Amaranthus* species. We identified a total of 445 CYP genes in three *Amaranthus* species, which were grouped into A-types and non-A-types, and further divided into nine clans and 39 families. The numbers of CYP clans and families vary among different plant species; for examples, Arabidopsis has 9 clans and 47 families (Bak et al. 2011), while soybean has 10 clans and 48 families (Khatri et al. 2022). Among the identified clans, clan 71 contains most of the *Amaranthus* CYP genes, whereas clans 74, 97, 710, 711, and 727 have only a few genes, reflecting their ancient origin and conserved nature (Bak et al. 2011). The presence of single-family clans 74, 97, and 710 in chlorophytes and charophytes indicates their deep evolutionary roots. Additionally, the emergence of the multi-family clan 71 and the single-family clan 711 is linked to the origin of mosses, with clan 71 expanding across various extant plant genomes (Durst and Nelson 1995; Nelson and Werck-Reichhart 2011; Hansen et al. 2021), 2021). The critical role of clan 71 in plant metabolism, stress adaptation, defense, and development may explain its large number of CYP genes.

Findings from this study revealed significant gene expansion (i.e., the presence of multiple copies of the same gene) and conservation (i.e., the same gene appearing across multiple species) at the family and subfamily levels. Notably, CYP76, CYP96, and CYP81 families show expansion across the three *Amaranthus* species. Furthermore, in *A. hybridus* and *A. retroflexus*, CYP71 and CYP72 families were also expanded. Some CYP families were unique to certain species; for example, CYP703 was found exclusively in *A. retroflexus*, and CYP712 was specific to *A. palmeri* (Supplemental Fig. S7A, S7B). The expansion and conservation of CYPs is a common phenomenon, and is likely influenced by the species’ habitat and lifestyle (Padayachee et al. 2020). Gain and loss of clans and families in *Amaranthus* species, consistent with findings from several other studies, were also observed (Helliwell et al. 2001; Nelson et al. 2004; Chhun et al. 2007; Li and Wei 2020).

Molin et al. (2020) reported the first complete assembly of a 399 Kbp eccDNA molecule that included *EPSPS* in *A. palmeri*. This eccDNA had little synteny when compared to reference genomes of closely related *Amaranthus* that were available at the time. Similar approaches using the new Hap1 genome assembly of *A. palmeri* indicate similar limited synteny between the canonical *EPSPS*-containing eccDNA and the Hap1 genome (Supplemental Fig. 8A). However, when alignment filters are relaxed, most of the eccDNA sequence does find a match in the genome (Supplemental Fig. 8B). Therefore, we hypothesize that the eccDNA is built from highly repetitive sequence from the *A. palmeri* genome that has undergone significant duplication and rearrangement. Regions of the eccDNA that lack matches even under lax filtration may be regions that were not assembled and included in the reference genome for one reason or another.

Here, we report a contiguously assembled circular sequence containing *EPSPS* that is highly syntenic to the canonical *EPSPS*-containing eccDNA molecule from a glyphosate-resistant plant from Georgia. Structural variants in this new sequence compared to the previously reported sequence are conserved across newly sequenced individuals in our study, suggesting our sequence to be more representative of the eccDNA molecule that has proliferated throughout the world. Our analysis finds no evidence of *EPSPS* being inserted into diverse eccDNA molecules (Fig. 5A). In fact, the lack of sequence divergence in *EPSPS* sequence contained within eccDNAs between geographically diverse accessions suggests a single duplication event of *EPSPS* into an eccDNA that has since proliferated throughout the world without generating almost any sequence diversity (Fig. 5B; Molin et al. 2020a). These results highlight the rarity of the generation of *de novo* insertions of herbicide target site genes into eccDNAs and the need for containment of recently identified herbicide-resistant weed populations.

In sum, our study highlights genomic features in *Amaranthus* species. We revealed candidate genes that are likely involved in sex determination in *A. palmeri*. The development of a working transformation protocol for *A. palmeri* would facilitate the functional validation of these candidates. While gene order was fairly conserved across the assemblies of *Amaranthus* species, chromosomal fusions and other structural rearrangement events contribute to the evolution of chromosomes in the genus. Our study also identified and classified cytochrome P450 genes in each studied species, facilitating investigations of the role of these genes in non-target site herbicide resistance and other phenotypes. Finally, we explore the genomic origins of eccDNA in *A. palmeri* and note conservation of amplified *EPSPS* sequence across diverse accessions. Moving forward, the genomic resources provided here will be valuable for understanding the mechanistic processes involved in herbicide resistance and in furthering the study of adaptive trait evolution within the genus.

## Materials and methods

### Plant material and growth conditions

The plant materials sequenced in this study are publicly available with the USDA Germplasm Resources Information Network (GRIN). *Amaranthus palmeri* and *A. retroflexus* seeds are available under the accession numbers PI 632235 and PI 572263, while the seeds of *A. hybridus* are from a population (FT-21605-14) that was sequenced and assembled in Montgomery et al. (2020). Seeds from the three species were sown separately in pots filled with premoistened soil (Lambert LM-GPS), and bottom irrigated. Seedlings were transplanted at ∼5 cm height into 16-cm pots (America Clay Works I-A650MP) filled with the same soil. Plants were grown at a temperature of 25/20 C and a photoperiod of 16 h/8 h (light/dark) regimes in greenhouses at Colorado State University. Tissue collection, DNA and RNA extractions, and library preparation have been previously described (Raiyemo et al. 2024). Library preparation and sequencing was performed at the Genome Center of Excellence at Corteva Agriscience.

### Genome sequencing, assembly, and annotation

The protocols adopted in the sequencing, assembly, and annotation of *A. palmeri*, *A. retroflexus*, and *A. hybridus* genomes were fully described in Raiyemo et al. (2024). Briefly, high molecular weight DNA was isolated from frozen leaf tissue of *A. palmeri* and *A. retroflexus* using Nucleobond HMW DNA kit. The isolated DNA was sheared and used to construct a SMRTBell HiFi library, followed by size selection. The 15 – 20 Kbp size selected fragments were then sequenced on PacBio Sequel Ile to an average depth of ∼50X the expected diploid genome size. Nuclei were extracted from young leaf tissue of each of the three species for Bionano optical mapping. Leaf tissue was fixed in 2% formaldehyde, chopped, homogenized, and filtered through cell strainers according to manufacturer’s protocol using the Prep™ Plant Tissue DNA Isolation Kit (Bionano Genomics, San Diego CA). The resulting nuclei were resuspended and cleaned from debris and other solids using a series of low-speed centrifugations (100 x g) followed by centrifugation at 2,500 x g prior to agarose embedding and purification. No gradient centrifugation was used. The Bionano Direct Label and Stain (DLS) protocol was performed as previously described (Hufford et al. 2021) and the labeled stained samples loaded into a single Bionano chip in a Saphyr system. The assembly of the genome maps and construction of hybrid scaffolds were performed using Bionano Access version 1.7 and Tools 3.7 on a local Bionano Computer server. Additionally, DNA was isolated from frozen leaf tissue of each of the three species, and chromatin was crosslinked with formaldehyde for Hi-C library preparation and sequencing on Illumina NovaSeq 6000.

*Amaranthus palmeri* and *A. retroflexus* genomes were assembled independently using Hifiasm v0.16.1 (Cheng et al. 2021). The resulting contig assemblies were then filtered and aligned to Bionano maps using the Bionano Solve software (v3.7_03302022). *Amaranthus hybridus* assembled contigs reported in Montgomery et al. (2020) were also aligned to a newly generated Bionano map. The hybrid scaffolds obtained were manually curated, and the matching pairs were confirmed to correspond to two haplotypes for each of the species. Sequence polishing was performed on the hybrid scaffolds by aligning PacBio HiFi corrected reads to the assemblies using minimap2 v2-2.24-r1122 (Li 2018). Hybrid scaffolds not assigned to any of the pseudomolecules were assigned to Chr00 in each species/haplotype. Hi-C data for each of the species were then used to validate the assembly of pseudomolecules using Juicer v0.7.0 (Durand et al. 2016b). To confirm that the haplotypes were resolved for *A. palmeri* and corresponded to the expected number of pseudomolecules per haplotype, the respective Hi-C map contacts were visualized using Juicebox (Durand et al. 2016a). Only one haplotype representative was reported for *A. retroflexus* and *A. hybridus*.

Annotation was performed in two steps: gene models were predicted, and functions were attributed. RepeatModeler v2.0.2, RepeatMasker v4.1.2 (Flynn et al. 2020), and BEDTools v2.30.0 (Quinlan and Hall 2010) were employed to generate softmasked genomes. Iso-seq PacBio reads were then aligned to the softmasked genomes using pbmm2 v1.10.0 (https://github.com/PacificBiosciences/pbmm2), and transcripts with minimum alignment coverage of 90% and minimum alignment identity of 95% were collapsed to unique isoforms utilizing IsoSeq3 v3.8.2 (https://github.com/PacificBiosciences/IsoSeq). The final gene model predictions were performed with MakerP v1.0 (Campbell et al. 2014) using the softmasked genome, the collapsed gene predictions from IsoSeq3, the classified repeat consensus libraries from RepeatModeler, and protein sequences from *Amaranthus* species available on NCBI. Filtering steps were performed with custom scripts to remove small proteins. Functional annotation combined several strategies to predict protein localization, function, and homology, through the use of tools including MultiLoc2 v1.0 (Blum et al. 2009), InterProScan5 v5.47 (Jones et al. 2014) and Uniprot, MMSeqs2 v4.1 (Steinegger and Söding 2017), and databases Uniref50, KEGG Orthology (KO), and NCBI. The genome annotation file includes relevant functional information in the notes column for each gene including the InterPro IDs, GO IDs, and the closest known annotated protein. Further details about genome annotation are described in Raiyemo et al. (2024). Transfer RNA (tRNA) genes were predicted using tRNAscan-SE v2.0.12 (Chan et al. 2021), which is part of the Maker pipeline.

Assembly quality and completeness were determined using the embryophyta_odb10 library within Benchmarking Universal Single-Copy Orthologs (BUSCOs) v4.0.2 (Simão et al. 2015). Additional assembly characteristics were computed using “agat_sp_statistics.pl” from AGAT Toolkit v1.0.0 (Dainat 2022) and “bbstats.sh” from BBTools (Bushnell 2014). The quality of intergenic and repetitive sequence space of all assemblies was also accessed using LTR assembly index (LAI) from the LTR retriever pipeline (Ou et al. 2018; Ou and Jiang 2018). Putative centromeric repeats in all assemblies were identified by searching the assemblies for centromeric repeats using CentroMiner (Lin et al. 2023). Telomeric repeats were identified by searching the telomere repeat motif (TTTAGGG)_4_ against all assemblies using BLASTN (Camacho et al. 2009).

### Sex determination GWAS in *A. palmeri*

An *A. palmeri* RAD-seq dataset (192 females and 192 males) that was previously used to develop sex specific markers by Montgomery et al. (2019) was used for genome-wide association analysis. The single-end raw reads data were demultiplexed, and cleaned using the “process_radtags” command in Stacks version 2.65 (Rochette et al. 2019), followed by mapping each sample to the Hap1 assembly of *A. palmeri* using BWA-MEM v0.7.17 (Li 2013). The “gstacks” command from Stacks was then used to build loci from the aligned reads. We grouped the individuals by sex, and then ran the “populations” command to filter out rare alleles (--min-maf 0.05). We then accessed the level of missingness in the two data groups using VCFtools v0.1.16 (--missing-indv) following a previous approach (Cerca et al. 2021). We removed individuals that had greater than 60% missing data, and then ran the “populations” command once again on the filtered samples utilizing additional filtering criteria (--min-maf 0.05, -p 2, -r 0.4). Using the set of variants obtained, we performed a principal component analysis with PLINK v1.90b7 (Chang et al. 2015), and then carried out a genome-wide association analysis with BLINK-C (Huang et al. 2019) using the first five principal components from PLINK to account for population stratification and familial relatedness. Sex of the individuals were used as binary input (phenotypes) in the analysis. Calculation of f-statistics (*F_ST_*) was carried out using VCFtools. Manhattan and *F_ST_* plots were generated using the CMplot package (Yin et al. 2021) in R.

### Identification of previous sex markers and genes in the *A. palmeri* assembly

A search of the previously reported primer sets, PAMS-940 (Montgomery et al. 2019) and JM940 (Montgomery et al. 2021), that amplified male-specific regions of *A. palmeri* was carried out against both haplotype assemblies with BLASTN (Camacho et al. 2009) using the -task blastn-short parameter. Reciprocal best hit (RBH) searches between genes on the previously identified scaffold 20 and the haplotype assemblies, as well as between the two haplotype assemblies, were carried out using MMseqs2 (Steinegger and Söding 2017).

### Synteny and intergenomic analyses

MCscan-python version (Tang et al. 2008) from JCVI utility libraries v1.1.22 was used to identify reciprocal best hit (RBH) and collinear gene blocks between haplotype 1 and 2 of *A. palmeri* genome assemblies. MCScanX (Wang et al. 2012) was also used to investigate duplicated genes within each of the *A. palmeri* haplotypes and in *A. retroflexus*, and *A. hybridus* genome assemblies.

To plot the duplicated genes with circos (Krzywinski et al. 2009), the output collinearity files from MCScanX were converted to “.anchors” format using “jcvi.compara.synteny” from JCVI. Sequence alignment between the two haplotypes of *A. palmeri* assemblies was carried out using Minimap2 v2.24-r1122 (Li 2021), and structural rearrangements between the two haplotypes were further refined using SyRI (Goel et al. 2019). Alignment between Hap1 and a previous draft assembly of a male *A. palmeri*, which contained male-specific regions on scaffold 20, was carried out using Minimap2. The alignments were visualized as dotplots using D-Genies (Cabanettes and Klopp 2018). Syntenic orthologues among available chromosome-level *Amaranthus* species; three from this study and four (*A. hypochondriacus*, *A. cruentus*, *A. tricolor*, and *A. tuberculatus*) from previous studies were identified and visualized using GENESPACE v1.2.3 (Lovell et al. 2022).

### Transcriptome profiling and gene ontology (GO) enrichment analysis

Quality control (QC) assessed and adapter trimmed mRNA reads for three tissue types (mature flowers, shoot apical meristem and floral meristems) from four biological replicates of each sex from Bobadilla et al. (2023) were mapped to Hap1 of the *A. palmeri* genome assembly using STAR v2.7.10b (Dobin et al. 2013). Two replicates from the mature female flower category were removed from downstream analyses as in the previous study due to low mapping quality (61.89% and 31.45% uniquely mapped reads for replicate 2 and 4, respectively). Read counting was then carried out using featureCounts v2.0.6 from the subread package (Liao et al. 2014). Genes with a CPM value of less than 1 in at least 2 samples were filtered out and the counts were normalized using the TMM normalization in edgeR (Robinson et al. 2009). The count normalization was then followed by the analyses of differential expression (DE) between male and female individuals for each tissue type in edgeR. Genes were assessed as differentially expressed based on FDR < 0.05 and Log_2_FC > 1.2 thresholds using the ‘*glmTreat*’ function within edgeR. Heatmaps were constructed with pheatmap package after a log2-transformed normalization of read counts in DESeq2 (Love et al. 2014). The counts were filtered to retain only genes that were differentially expressed from the previous analysis prior to the heatmap construction.

The translated protein-coding sequences (CDS) of Hap1 were assigned GO annotations using eggNOG-mapper v2.1.12 (Cantalapiedra et al. 2021). GO term enrichment analysis was then carried out using topGO with nodeSize = 10. The enrichment test was performed using Fisher’s exact test and the “elim” algorithm. GO terms were assessed as significantly enriched at the default *p* < 0.01 threshold. The enrichment plot was generated using ggplot2 (Wickham et al. 2019) in R.

### Coverage analysis

To determine the conservation and specificity of genes within the MSY region of the *A. palmeri* Hap1 genome assembly, short reads of male and female *A. watsonii* individuals from previous studies (Raiyemo et al. 2023; Raiyemo and Tranel 2023) were mapped to the *A. palmeri* Hap1 genome using BWA-MEM v0.7.5 (Li 2013). SAMtools v1.14 (Danecek et al. 2021) fixmate was used to fill mate coordinates and insert size fields, and duplicates were marked with Picard v2.26.9 (https://broadinstitute.github.io/picard/). The alignment files were then filtered for uniquely mapped reads using SAMtools and “grep” to remove unmapped reads (-F 4), reads with MAPQ < 5, alternative hits (tag XA:Z), and supplementary alignments (tag SA:Z). Both the unfiltered and filtered reads for the male and female individuals were visualized in IGV (Thorvaldsdóttir et al. 2013).

### Cytochrome P450 identification, naming, and comparative analyses

To identify CYP genes in the annotated protein sequences of *A. palmeri*, *A. hybridus*, and *A. retroflexus*, “grep” was used to identify genes in the annotation file associated with the InterPro codes “IPR001128” and “IPR036396”. All identified protein candidates were blasted against all named plant CYPs (http://drnelson.uthsc.edu/CytochromeP450.html) and were submitted to the Standardized Cytochrome Nomenclature Committee (https://drnelson.uthsc.edu/CytochromeP450.html) to ensure consistent naming conventions.

Briefly, sequences demonstrating >40% identity were grouped in the corresponding family as the named homologous proteins, while sequences displaying >55% identity were categorized into the same subfamily. Proteins containing less than 350 amino acids were considered fragmented annotations and were discarded.

The amino acid sequences of all full-length CYP proteins from *A. palmeri, A. retroflexus* and *A. hybridus* were aligned using Clustal W (Chenna et al. 2003). A neighbor-joining (NJ) phylogenetic tree was constructed using MEGAX with 1000 bootstrap replications (Kumar et al. 2018). The resulting phylogenetic tree was visualized using the ITOL9 web server (https://itol.embl.de/). MCScanX (Wang et al. 2012) was used to identify the collinearity of CYP genes in the three species. TBtools (Chen et al. 2020) was then employed to map the collinear gene pairs of the CYPs.

### Origin of extrachromosomal DNA (eccDNA)

Palmer amaranth seeds were grown under the same conditions outlined for whole genome sequencing. Accessions included glyphosate-resistant populations from Georgia, Tennessee, Arizona, and Brazil as well as glyphosate-susceptible populations from Georgia, Arizona, Kansas, and USDA-S, a GRIN accession (PI 632235) originally collected in 1986 from Arizona that was also used to build the reference genome reported here. One plant per accession of unknown sex was selected for whole genome sequencing with PacBio HiFi technology as outlined above for genome assembly. Each sample was sequenced to a depth of at least 35X according to the haploid genome size of 410 Mbp.

Resequencing of eight plants yielded between 15 and 24 Gbp of PacBio HiFi sequence per individual. The reads of each sequenced sample were assembled *de novo* using Hifiasm v0.19.8 (Cheng et al. 2021). Assemblies were reported for both haplotypes, but only the Hap1 assembly was retained per individual. The number of contigs obtained for each assembly ranged from 833 to 1299 contigs with N50 values ∼2 Mbp. CD-HIT v4.8.1 (Fu et al. 2012) was used to collapse contigs with >95% similarity. BLAST v2.10.1+ (Camacho et al. 2009) was used to identify the remaining contigs from each assembly with all eight exons of the *A. palmeri EPSPS* gene model (AmaPaChr07Ag122820 in the Hap1 annotation). The *EPSPS* gene models (including introns) were manually extracted from these contigs for all individuals. Contigs containing the genomic version of *EPSPS* were identified by aligning the sequence of *UDP-glucuronate* (AmaPaChr07Ag122840 in the Hap1 annotation), a single-copy gene near *EPSPS* on chromosome 7. This gene was not coduplicated with *EPSPS* in the eccDNA molecule, so it was only found coassembled with the genomic version of *EPSPS*. A phylogenetic tree was generated to infer the evolutionary history of the extracted *EPSPS* sequences along with *EPSPS* sequences from several closely related species, using several *Amaranthus* species as outgroups. The Maximum Likelihood method and Tamura 3-parameter model (Tamura 1992) were used in MEGA X (Kumar et al. 2018) for tree construction.

Minimap2 v2.26 (Li 2018) was used to identify regions of synteny (map quality=60, minimum alignment length=1000) between the canonical eccDNA harboring *EPSPS* (Molin et al. 2020b) and the Hap1 assembly of *A. palmeri*. Surviving alignments were plotted with Circos (Krzywinski et al. 2009). Nucmer, part of the MUMmer v4.0.0rc1 package (Marçais et al. 2018), was also used to identify regions of similarity between the eccDNA and Hap1 (minimum cluster length=25, use all anchor matches regardless of their uniqueness). Surviving alignments were plotted with mummerplot. Minimap2 (Li 2018) was also used to identify regions of synteny (map quality=60, minimum alignment length=1000) between *EPSPS*-containing contigs from the resequenced assemblies and the canonical eccDNA (Molin et al. 2020b). These alignments were plotted using minidot (https://github.com/thackl/minidot).

### Data access

Sequencing data analyzed in this study are available through the National Center for Biotechnology Information (NCBI) (*A. palmeri*, PRJNA1120405; *A. retroflexus*, PRJNA1120413; *A. hybridus*, PRJNA1125363; and *A. palmeri* resequencing, SRA accession SRR30167488-SRR30167495). Genome assemblies including FASTA files and accompanying GFF annotation files have been deposited on NCBI (*A. palmeri* Hap1; JBEFMX000000000, *A. palmeri* Hap2; JBEFMY000000000, *A. retroflexus*; JBEFNP000000000, and *A. hybridus*; JBELOC000000000), CoGe [*A. palmeri* Hap1 (68302), *A. palmeri* Hap2 (68303), *A. retroflexus* (68315), and *A. hybridus* (68316)], and the International Weed Genomics Consortium online database WeedPedia (https://www.weedgenomics.org/species/amaranthus-palmeri/, https://www.weedgenomics.org/species/amaranthus-retroflexus/, https://www.weedgenomics.org/species/amaranthus-hybridus/). A 395 Kbp circular contig containing *EPSPS* from the assembly of a glyphosate-resistant plant from Georgia has been deposited on NCBI (GenBank accession number: PQ096843).

## Acknowledgements

This work was supported by the International Weed Genomics Consortium with funding from Foundation for Food & Agriculture Research (FFAR grant number DSnew-0000000024), Bayer AG, Corteva Agriscience, Syngenta Ltd, BASF SE, and CropLife International (Global Herbicide Resistance Action Committee). Funding was also provided by the USDA National Institute of Food and Agriculture (Grant number 2022-67013-36142 to PJT).

## Author’s contributions

P.J.T., T.A.G., E.L.P., C.A.S., and D.R.N conceived the research study. J.S.M. and S.M. grew plants, harvested, and prepared samples for shipping to the sequencing center. V.L. and K.F. performed DNA and RNA extraction, PacBio HiFi, Bionano DLS, Hi-C, Iso-Seq sequencing, as well as genome assembly. E.L.P. and L.C. annotated the genomes and provided genome assembly metrics. D.A.R. carried out genome-wide association analysis, transposable elements analysis, and comparative genomics analyses. J.S.M. carried out the eccDNA analysis. F.A. and D.R.N. carried out the cytochrome P450 analysis. A.J.L. carried out the coverage analysis. D.A.R., J.S.M., L.C., and F.A. wrote the manuscript. All authors read, revised, and approved the manuscript.

